## Supplemental Table and Figures for "Age-related differences in immune dynamics during SARS-CoV-2 infection in rhesus macaques"

**Supplemental Table 1. Clinical signs in rhesus macaques inoculated with SARS-CoV-2.**

| Animal ID | Observed clinical signs | | | |
|---|---|---|---|---|
| | 1-3 dpi | 4-7 dpi | 8-14 dpi | 15-21 dpi |
| **RMO1** | Pale appearance, reduced activity, mildly depressed, tachypnea | Pale appearance, reduced activity, mildly depressed, ruffled fur, tachypnea, dyspnea | N/A | N/A |
| **RMO2** | Tachypnea, dyspnea | Slightly reduced appetite, nosebleed on 4 dpi, dyspnea | N/A | N/A |
| **RMO3** | Pale appearance, reduced activity, ruffled fur, dyspnea | Pale appearance, reduced activity, slightly ruffled fur, slightly reduced appetite, mildly dehydrated, dyspnea | N/A | N/A |
| **RMO4** | Ruffled fur, mildly dehydrated | Reduced activity, ruffled fur, severely reduced appetite, mildly dehydrated, dyspnea | N/A | N/A |
| **RMO5** | Pale appearance, hunched posture, reduced activity, mildly depressed, reduced appetite | Reduced activity, depressed, reduced appetite | Mildly depressed | Mildly depressed, reduced appetite |
| **RMO6** | Reduced appetite | Reduced appetite | Slightly reduced appetite | Slightly reduced appetite |
| **RMO7** | Ruffled fur, reduced appetite, tachypnea | Ruffled fur, severely reduced appetite, tachypnea | Ruffled fur, severely reduced appetite, tachypnea | Severely reduced appetite, tachypnea |
| **RMO8** | Ruffled fur, tachypnea | Ruffled fur, reduced appetite, tachypnea | Ruffled fur, reduced appetite, tachypnea | Tachypnea |
| **RMY1** | Slightly reduced appetite, tachypnea | Slightly reduced appetite | N/A | N/A |
| **RMY2** | No signs | Reduced appetite, serous nasal discharge on 6 dpi, dyspnea | N/A | N/A |
| **RMY3** | Dyspnea | Slightly reduced appetite, dyspnea | N/A | N/A |
| **RMY4** | Pale appearance, tachypnea, dyspnea | Slightly reduced appetite, dyspnea | N/A | N/A |
| **RMY5** | Slightly reduced appetite, dyspnea | Dyspnea | No signs | Slightly reduced appetite, recovered by 18 dpi |
| **RMY6** | Ruffled fur, reduced appetite, tachypnea | Ruffled fur, tachypnea, dyspnea | Dyspnea | Recovered |
| **RMY7** | Slightly reduced appetite, tachypnea | Slightly reduced appetite, tachypnea | Slightly reduced appetite, tachypnea | Slightly reduced appetite, tachypnea through 17 dpi |
| **RMY8** | Ruffled fur, reduced appetite | Hunched posture, ruffled fur, tachypnea, | Dyspnea | Recovered |

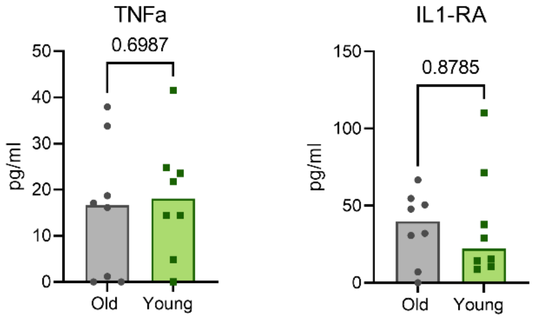


**Supplemental Figure 1. Baseline cytokine levels.** Expression of serum cytokines TNFα and IL1-RA in older and younger animals at 0 dpi. Points represent individual animals; bars represent the group median. P-values are calculated using a Mann-Whitney test.

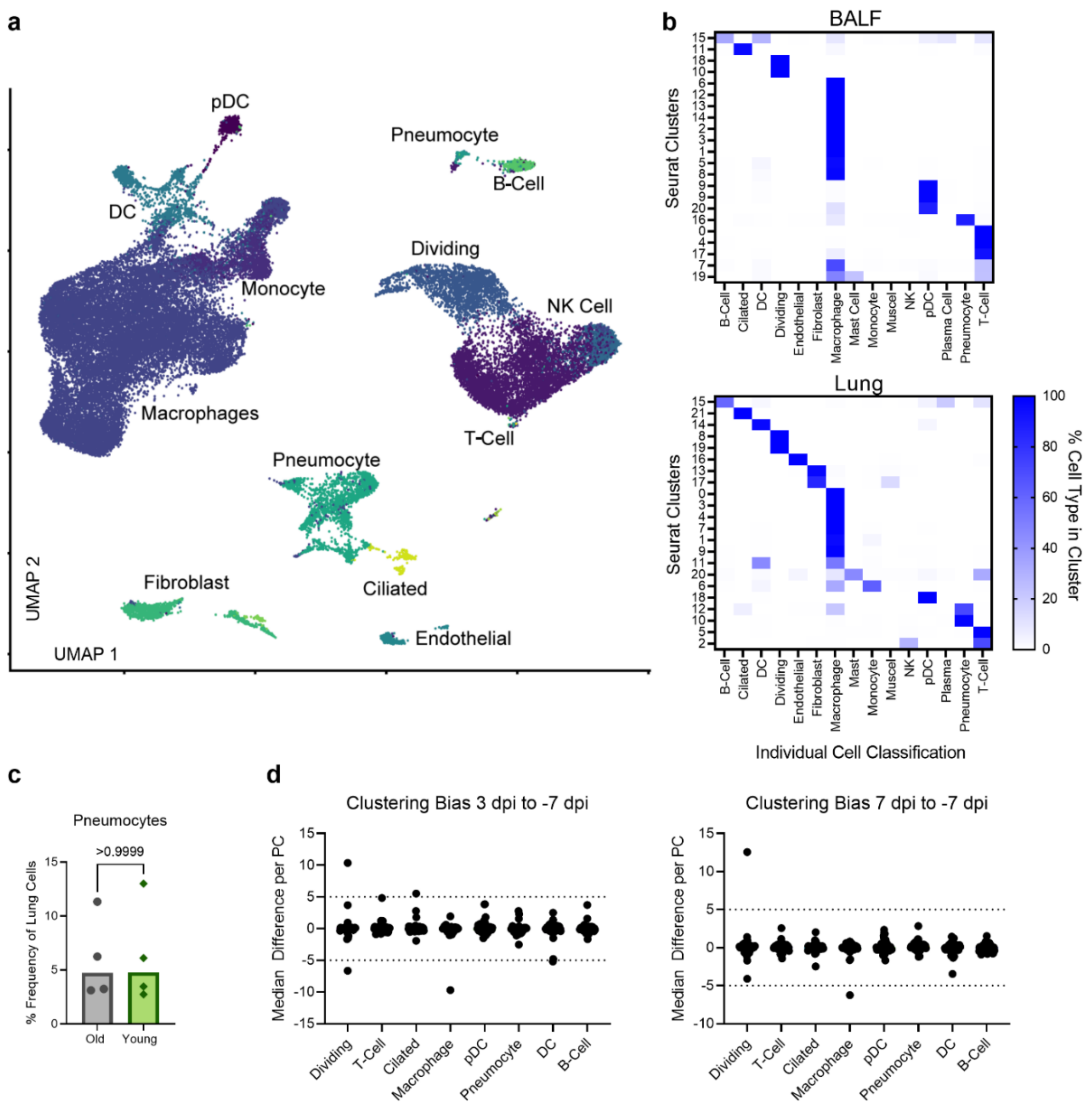


**Supplemental Figure 2. Single cell RNA sequencing of lung and cell type identification with clustering bias. (a)** UMAP projection from single cell RNA sequencing of lung sections at 7 dpi. Each point is an individual cell and is colored based on its determined individual cellular phenotype. Cell phenotypes are listed next to the clusters. **(b)** Comparison of computationally determined individual cell type identification with Seurat-determined clusters at a resolution of 0.8 in BALF (top) and lung samples (bottom). Columns represent individual cell types and rows the Seurat Clusters. The color represents the percentage of cells in a given Seurat cluster that was identified as a specific cell type with white being 0% and darker blue being 100%. **(c)** The percentage of total lung cells identified as pneumocytes in older compared to younger animals. P-value is calculated in a Mann-Whitney test. **(d)** Clustering bias in BALF single cell sequencing samples across the different cell types. Each point represents a principal component within the given cell type. The difference in the median value between where points from either 7 dpi (left) or 3 dpi (right) fall along a given principal component as compared to baseline on -7dpi is shown. Dashed lines at 5 and -5 represent the cutoffs for when the difference is sufficiently large to warrant further analysis.

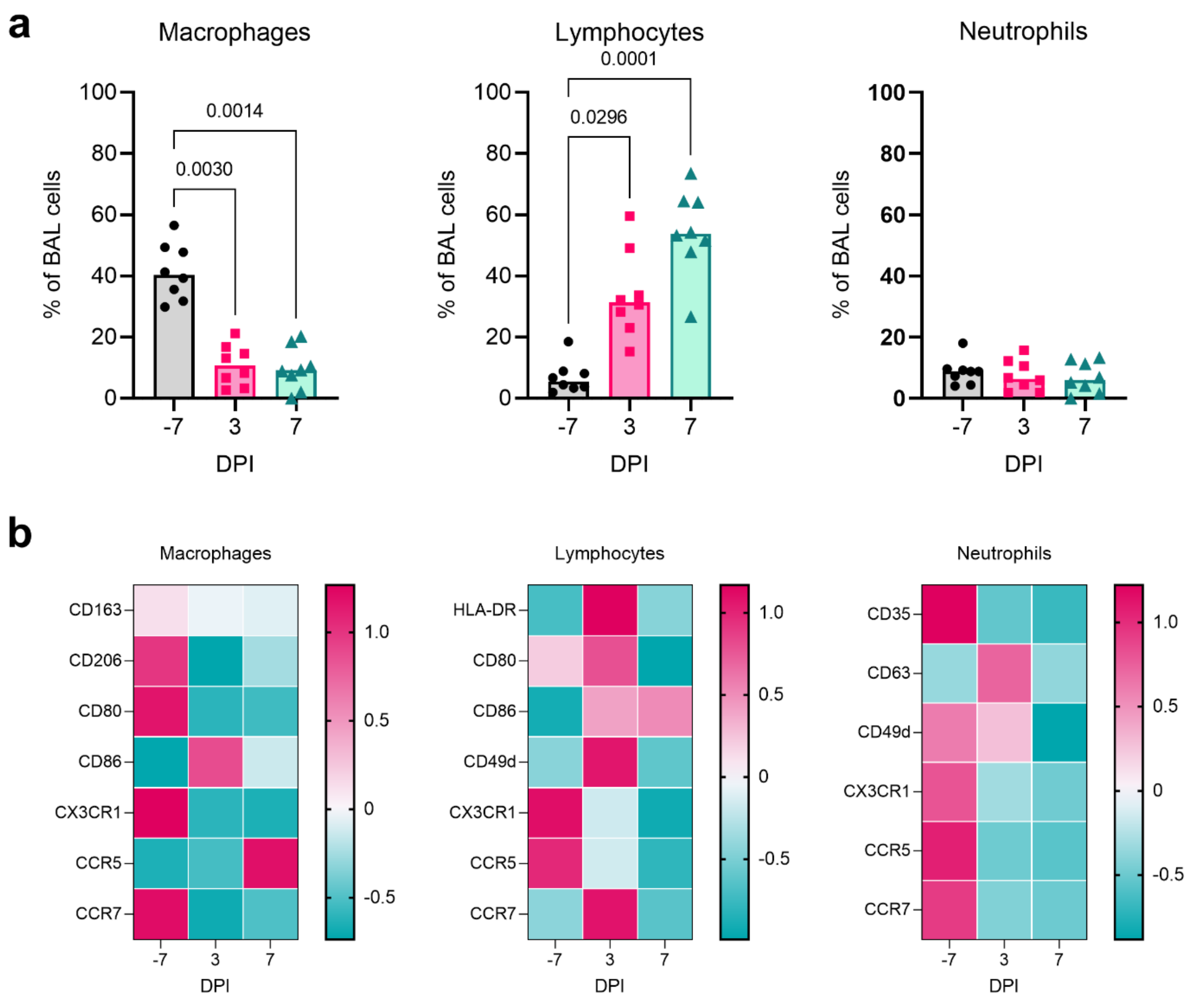


**Supplemental Figure 3. Flow cytometric validation of cell frequencies in BALF during SARS-CoV-2 infection. (a)** Frequency of cells in BALF as determined by flow cytometry for major cell lineages (macrophages, lymphocytes, and neutrophils) over time in one cohort of animals (group necropsied at 7 dpi). P-values are calculated in a 2-way ANOVA and only those < 0.1 are shown for clarity. **(b)** Heatmaps of the median fluorescence intensity (MFI) of different markers in macrophages (left), lymphocytes (middle), and neutrophils (right) over time. MFI data are z-normalized across the rows.

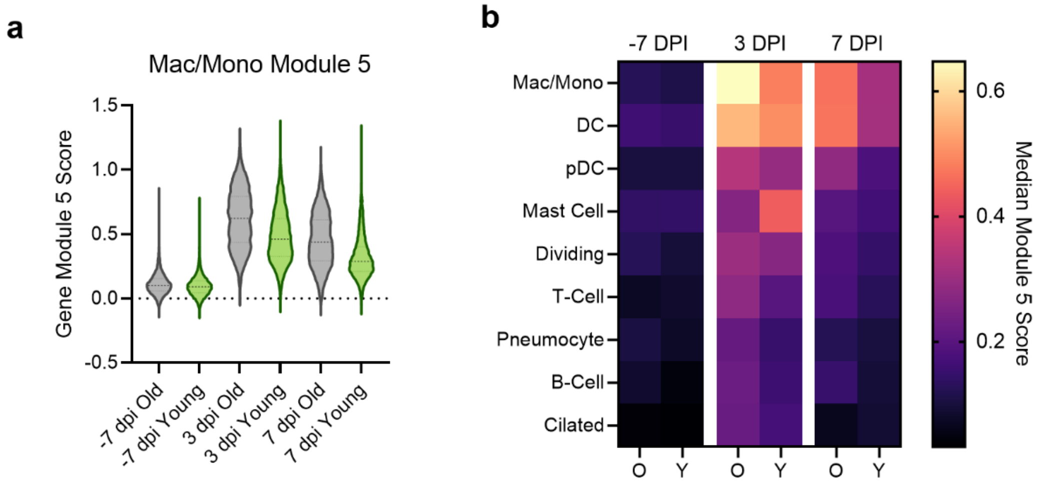


**Supplemental Figure 4. Gene Module 5 scores.** (Left) Macrophage Gene Module 5 score in BALF is shown for individual cells in older compared to younger animals over the time course (x-axis). Violin plots show the distribution; middle dashed line shows the median, and the smaller dashed lines represent the first quantiles. (Right) Heatmap of the macrophage gene module 5 median module score at pre-inoculation (-7 dpi), 3dpi, and 7 dpi across all cell types in the BALF. Each column pair represents the older compared to the younger animals.

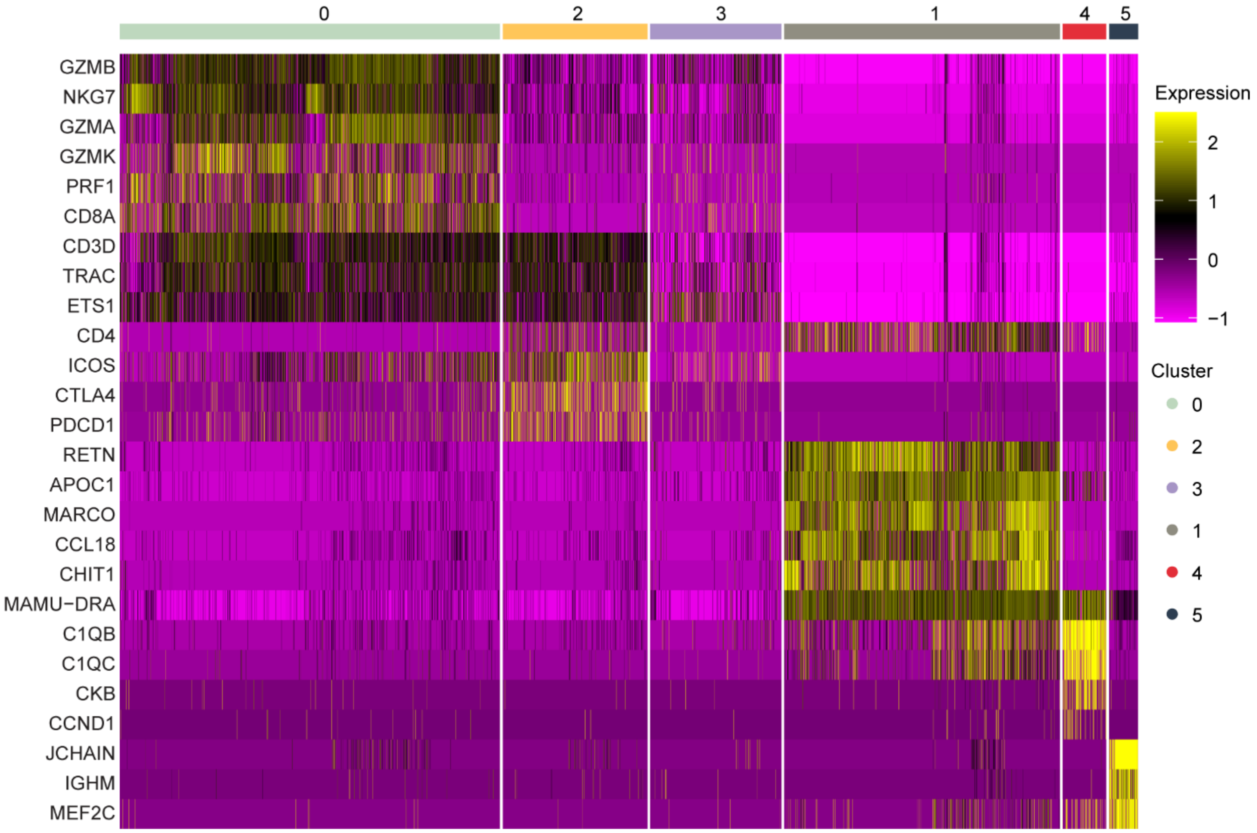


**Supplemental Figure 5. Dividing cell cluster gene expression of marker genes in BALF.** Marker gene expression in individual cells (columns) is shown from the BALF dividing cell cluster. The groups across the top are the Seurat clusters determined by a resolution of 0.2 as shown in Figure 4a. The colors represent the normalized expression of the gene in a given cell with purple representing low to no expression and yellow representing high expression values.

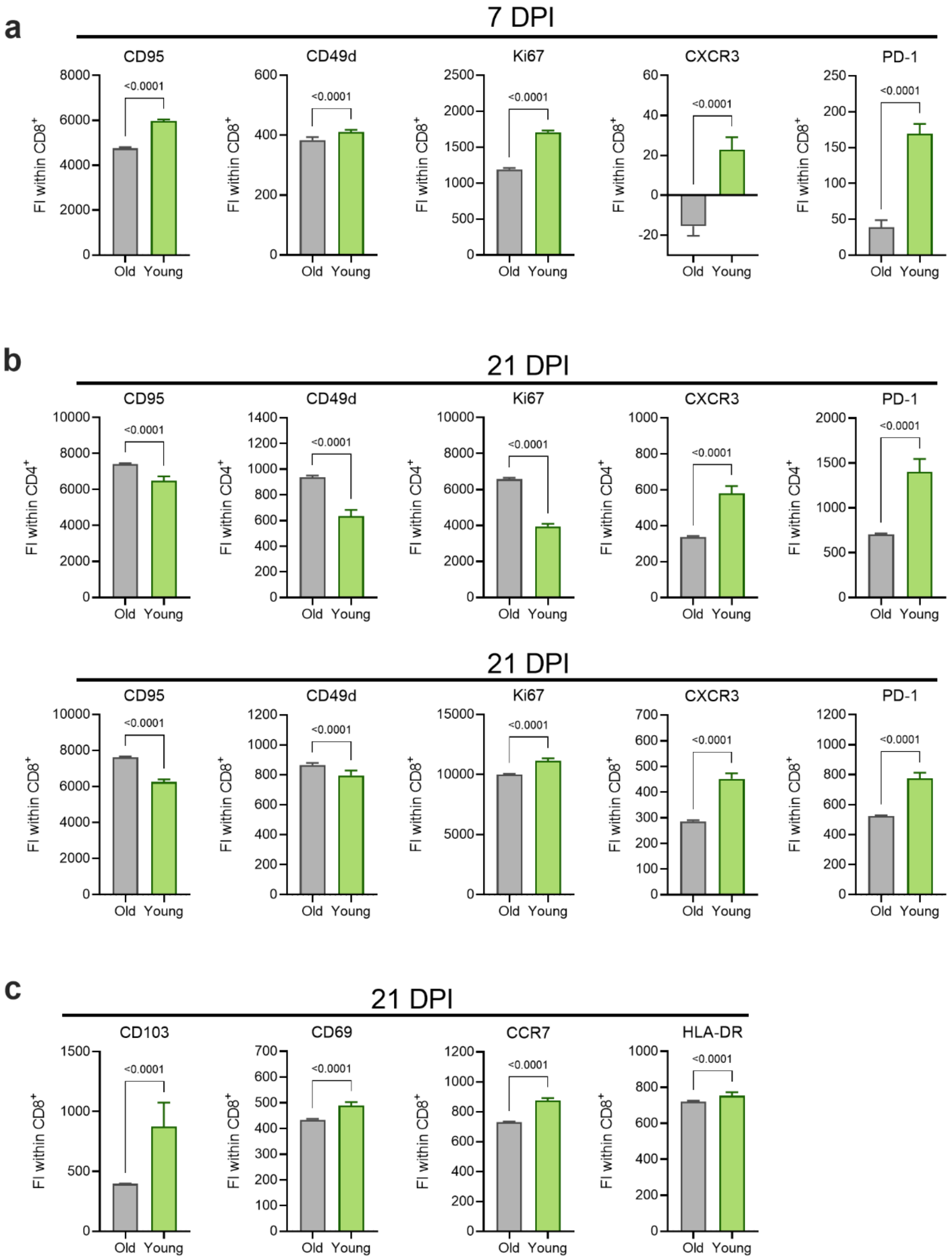


**Supplemental Figure 6. Age-associated expression patterns of markers associated with effector and memory function on CD4$^+$ and CD8$^+$ lung T-cells at 7 and 21 DPI.** The fluorescence intensities (FI) of selected markers stained by flow cytometry on CD8$^+$ T-cells at 7 dpi **(a)**, CD4$^+$ (top) and CD8$^+$ (bottom) T-cells at 21 dpi **(b)**, and CD8$^+$ T-cells at 21 dpi **(c)** were compared between older and younger rhesus macaques. Bars depict the median and 95% CI calculated across FI values generated for individual CD4$^+$ and CD8$^+$ T-cells; the data from these subsets were concatenated across animals within each cohort. P-values are calculated using Mann-Whitney tests.

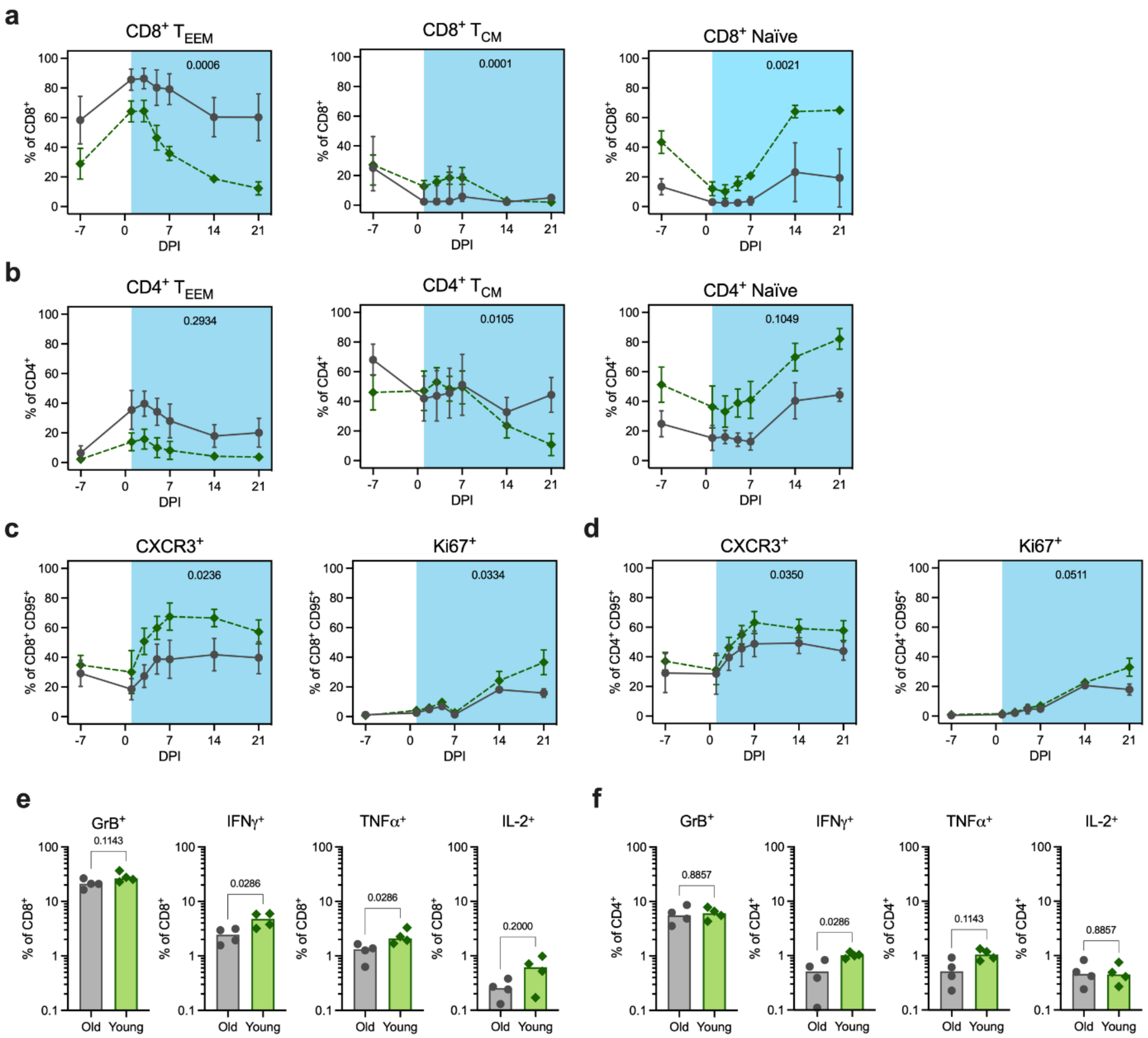


**Supplemental Figure 7. Circulating and splenic T-cell responses post inoculation.** Frequencies of effector and effector memory ($T_{EEM}$, left), central memory ($T_{CM}$, middle), and naïve (right) $CD8^+$ **(a)** and $CD4^+$ **(b)** T-cells are shown over time (dpi) as a percentage of total $CD8^+$ and $CD4^+$ T-cells, respectively. Frequencies of $CXCR3^+$ (left) and $Ki67^+$ (right) populations are shown over time (dpi) as a percentage of $CD8^+$ **(c)** and $CD4^+$ **(d)** non-naïve ($CD95^+$) T-cells. Data points represent the mean frequencies with SD (error bars) connected across timepoints for the older (grey circle, solid line) and younger (green diamond, dashed line) rhesus macaques. AUC of frequencies from 1 to 21 dpi were compared between the age cohorts by unpaired t-test and p-values are indicated on the graphs. The percentage of splenic $CD8^+$ **(e)** and $CD4^+$ **(f)** T-cells expressing (from left to right) Granzyme B (GrB), IFNγ, TNFα, and IL-2, after SAR-CoV-2 spike protein-peptide pool stimulation and intracellular cytokine staining. All bars depict group medians; p-values are the result of Mann-Whitney tests.

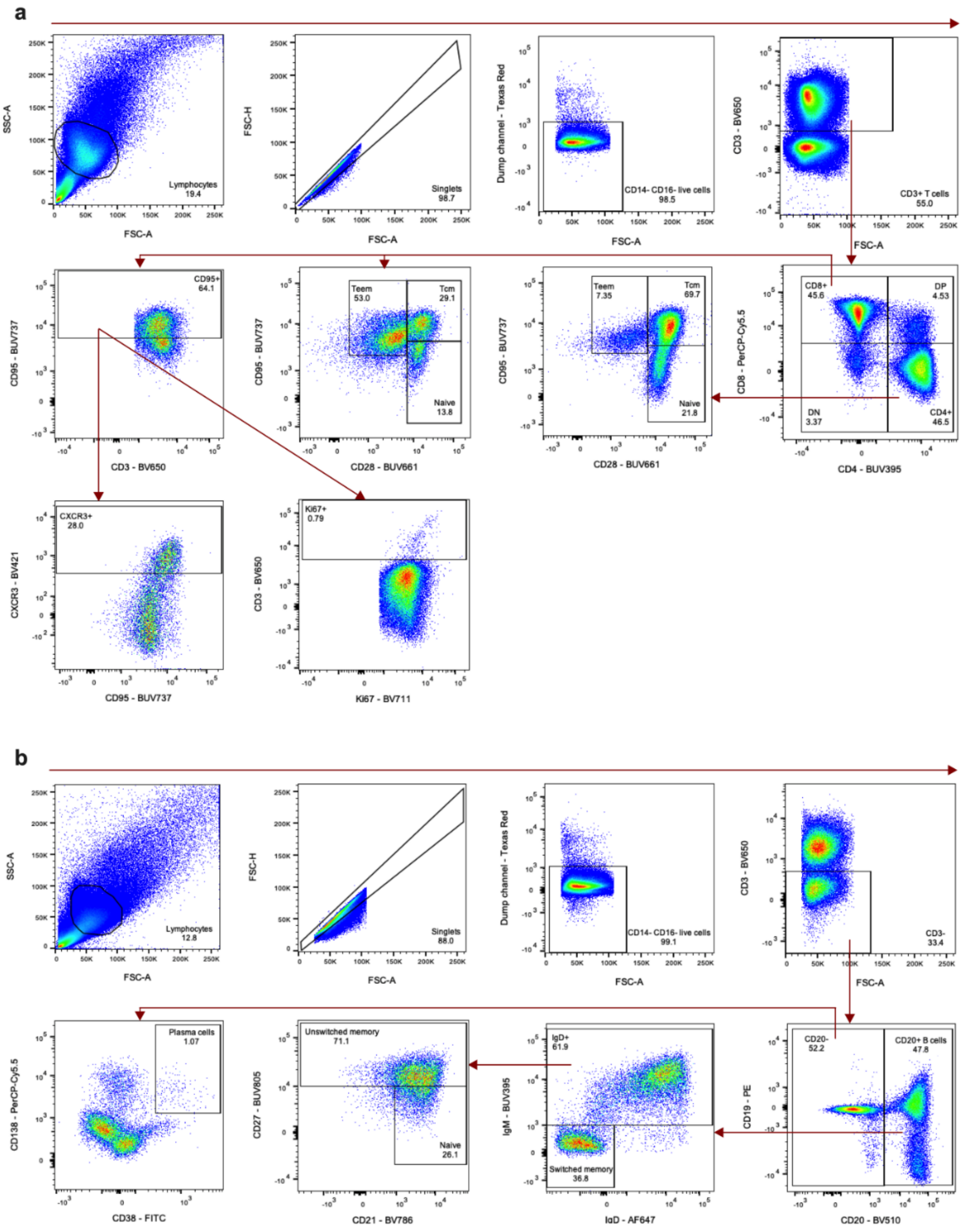


**Supplemental Figure 8. Flow cytometry gating strategy for characterization of T-cells and B-cells in PBMC samples.** All events were depicted using forward scatter area (FSC-A) and side scatter area (SSC-A) and lymphocytes were gated. Singlets were isolated using FSC-A and forward scatter height (FSC-H).

Dead cells, $CD14^{+}$, and $CD16^{+}$ cells were excluded by gating on negative cells in the dump channel. **(a)** T-cells were identified as $CD3^{+}$, and further classified as $CD4^{+}$ ($CD4^{+}$ $CD8^{-}$), $CD8^{+}$ ($CD4^{-}$ $CD8^{+}$), DP ($CD4^{+}$ $CD8^{lo}$), and DN ($CD4^{-}$ $CD8^{-}$). Memory subsets of $CD4^{+}$ and $CD8^{+}$ T-cells were characterized as follows: naïve ($CD28^{+}$ $CD95^{lo}$), central memory ($T_{CM}$, $CD28^{+}$ $CD95^{hi}$), or effector/effector memory ($T_{EEM}$, $CD28^{-}$ $CD95^{+}$). $CD8^{+}$ T-cells were additionally gated to isolate the total non-naïve ($CD95^{+}$) population; within this gate, $CXCR3^{+}$ and $Ki67^{+}$ cells were identified. **(b)** B-cells were classified as $CD3^{-}$ and $CD20^{+}$. Class-switched memory B-cells were further characterized as $IgD^{-}$ $IgM^{-}$. Within the $IgD^{+}$ gate, unswitched memory and naïve B-cells were identified as $CD27^{+}$ and $CD27^{-}$ $CD21^{+}$, respectively. Plasma cells were isolated within the $CD20^{-}$ gate and classified as $CD38^{+}$ $CD138^{+}$.

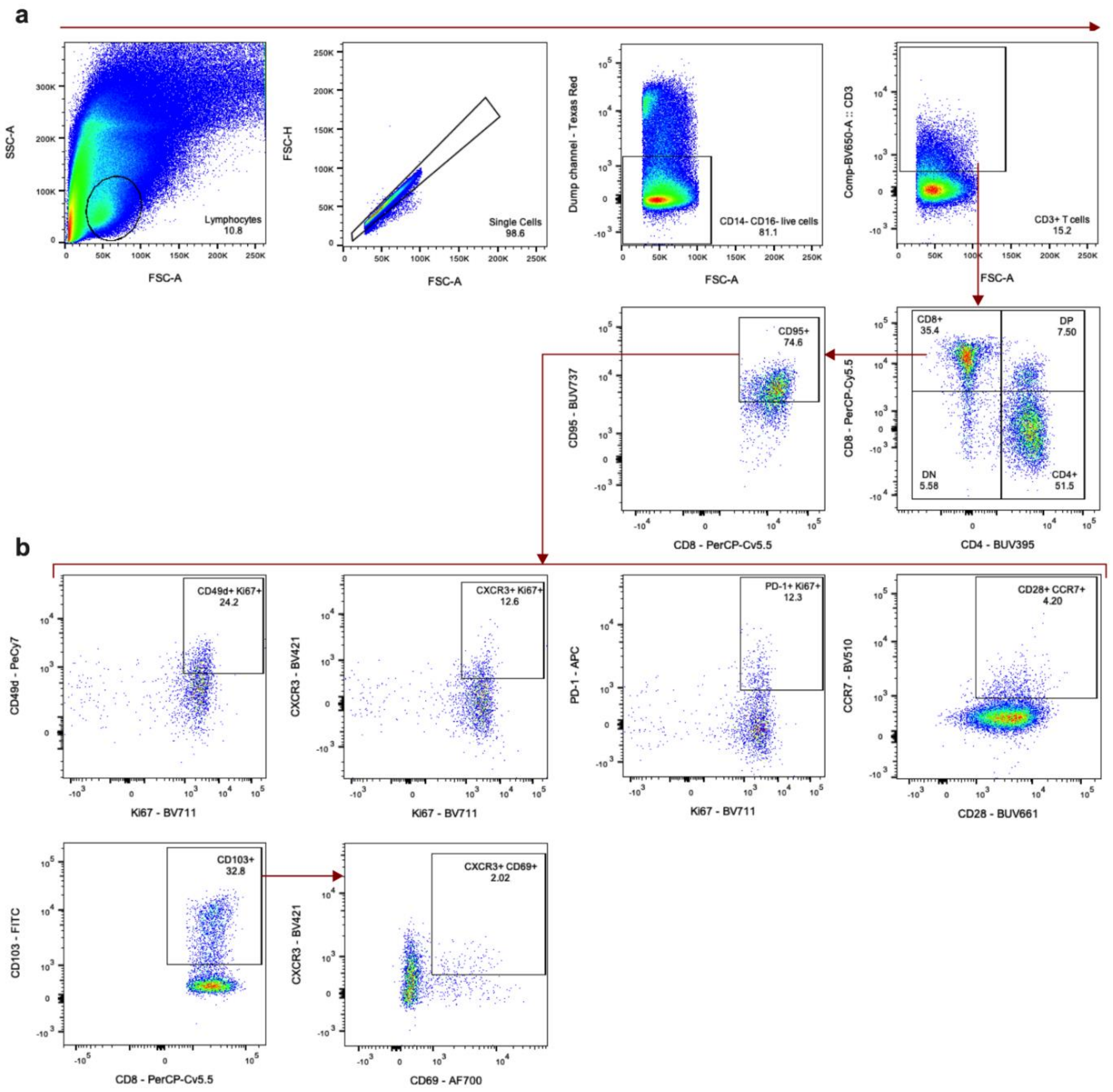


**Supplemental Figure 9. Flow cytometry gating strategy for characterization of T-cells in the lungs.** **(a)** Classification of T-cells and major T-cell subsets in the lungs was performed as described in Figure S8a. **(b)** Within the $CD8^{+}$ non-naïve ($CD95^{+}$) T-cell gate, the following populations were identified: $CD49d^{+}$ $Ki67^{+}$, $CXCR3^{+}$ $Ki67^{+}$, PD-$1^{+}$ $Ki67^{+}$, $CD28^{+}$ $CCR7^{+}$, and $CD103^{+}$. A sub-population of $CXCR3^{+}$ $CD69^{+}$ cells was further isolated within the $CD103^{+}$ gate.

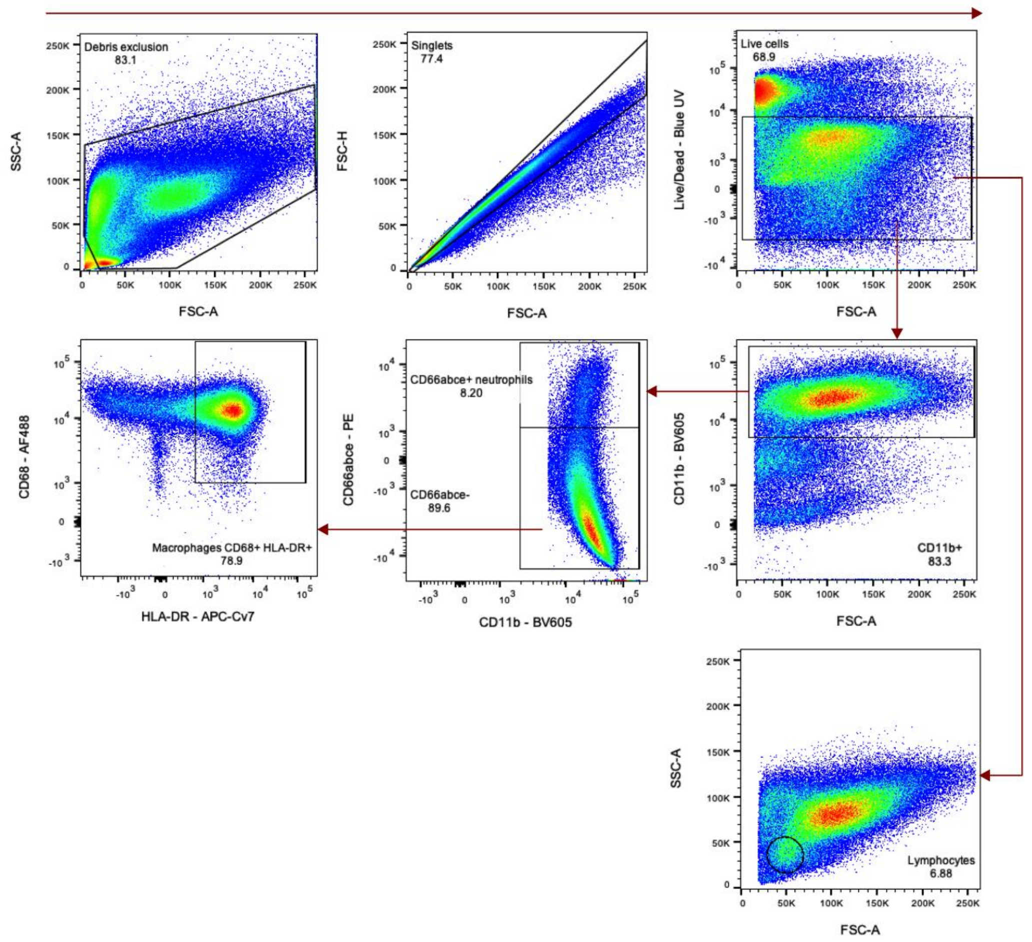


**Supplemental Figure 10. Flow cytometry gating strategy for cell type identification in BALF.** Gating to exclude debris was performed using FSC-A and SSC-A prior to doublet discrimination using FSC-A and FSC-H. Dead cells were excluding by gating on the population of cells negative for LIVE/DEAD staining. A $CD11b^{+}$ population was isolated; within this gate, neutrophils were classified as $CD66abce^{+}$. Macrophages were identified by gating on $CD66abce^{-}$ cells followed by $CD68^{+}$ $HLA\text{-}DR^{+}$ cells. Lymphocytes were classified as $FSC\text{-}A^{lo}$ and $SSC\text{-}A^{lo}$ within the live cell gate.

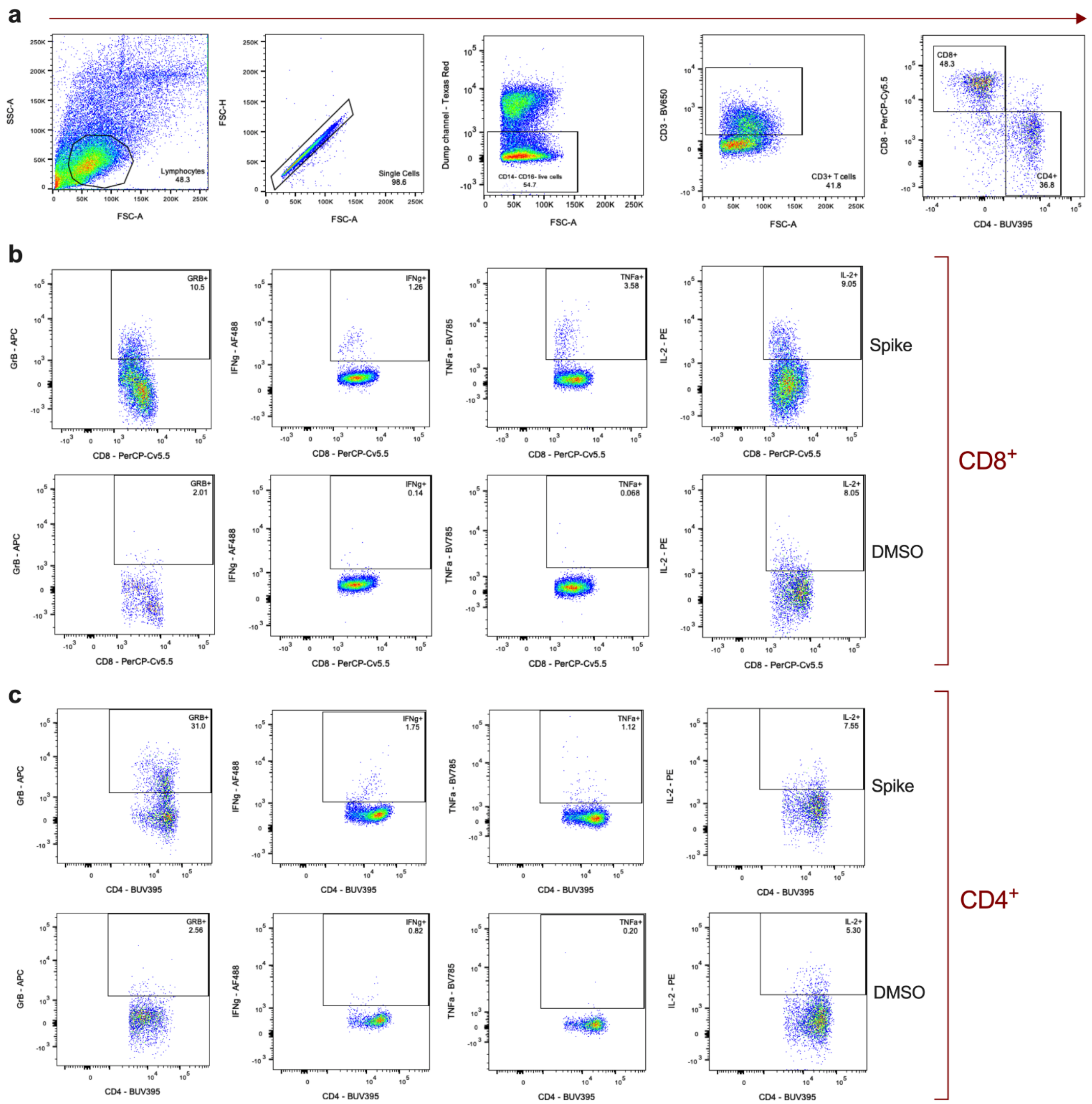


**Supplemental Figure 11. Flow cytometry gating strategy for intracellular cytokine staining of splenocytes. (a)** Identification of total, CD8$^+$, and CD4$^+$ T-cells was performed as described in Figure S8a. Gating to isolate single-cytokine positive populations (GrB, IFN$\gamma$, TNF$\alpha$, and IL-2) within CD8$^+$ **(b)** and CD4$^+$ **(c)** T-cell subsets is indicated. An example of a sample stimulated with the spike (S1) protein-peptide pool (top) is shown along with the corresponding DMSO-treated control (bottom) for each cytokine.
